# Deepening sleep using an EEG wearable featuring modeling-based closed-loop neurostimulation

**DOI:** 10.1101/2025.03.09.642252

**Authors:** Stefan L. Jongejan, Anna C. van der Heijden, Lucia M. Talamini

## Abstract

**Objective:** Closed-loop neurostimulation (CLNS) during slow-wave sleep (SWS) has been shown to enhance slow-wave activity, predominantly using laboratory equipment. To further advance CLNS research and its potential applications, there is a need for user-friendly EEG wearable equipment featuring CLNS that can support long-term CLNS studies in both clinical and home settings.

**Approach:** Here we evaluate whether modeling-based CLNS (M-CLNS) with acoustic stimulation of slow oscillations (SOs) can be effectively implemented using a self-applicable EEG headband with forehead electrodes. We assess the performance of M-CLNS stimulus targeting over the EEG headband together with short-term, stimulus-locked, and more enduring enhancement of SWS. We validate our results against simultaneous recordings obtained using gold-standard laboratory PSG equipment.

**Main results:** Our findings demonstrate that the SO phase can be reliably assessed and accurately targeted using M-CLNS with the EEG headband. We show an immediate enhancement of SO dynamics in the short-term, as well as an enduring increase in power spectral density across SO and delta frequencies (0.75 – 5 Hz) across SWS. These results are in line with previous studies using M-CLNS with laboratory equipment.

**Significance:** These findings demonstrate that M-CLNS with acoustic stimulation can successfully be applied using an EEG headband on the forehead to target SOs, leading to both immediate and enduring enhancements of SWS. In conclusion, M-CLNS in a self-applicable EEG headband may offer a promising tool for portable and non-invasive enhancement of SWS, with future potential for clinical and home-based applications.

## 1. Introduction

Poor sleep quality has become a widespread concern in modern societies, affecting a substantial portion of the population. Over a third of adults in industrialized countries report poor sleep quality, with estimates reaching up to 60% in some regions [1-4]. Both the World Health Organization and the US Centers for Disease Control have declared sleep problems a public health epidemic. The impact of this epidemic is far-reaching, as sleep harbors essential growth and recovery processes that sustain physical and mental health. Accordingly, poor sleep is a significant cause of morbidity, mortality, accidents and work absenteeism, with a tremendous societal and economic cost.

The aspect of sleep that is critical for growth and recovery functions is deep sleep, also termed slow wave sleep (SWS). SWS has been shown to support the immune response [5,6], build-up of energy molecules [7], cardiovascular health [8], glucose homeostasis [9] and brain waste clearance [10].

A decline of SWS has been linked to a large variety of disorders. These include most sleep disorder including insomnia [11,12] and sleep apnea [13], many psychiatric conditions, like post-traumatic stress disorder [14] and depression [15], neurodegenerative diseases such as Alzheimer’s [16] and Parkinson’s disease [17], and somatic syndromes such as chronic fatigue syndrome [12]. Additionally, SWS naturally declines with age, affecting the globally increasing aging population [18] and contributing to the prevalence of SWS-related health concerns.

Despite this wide representation of SWS disturbance, available treatments are few. Sleeping problems are mostly addressed with pharmacological interventions, which are not directly targeting SWS disturbance. These interventions mostly add light sleep and suppress REM sleep [19], while minimally affecting SWS. In addition, these pharmacological interventions have other important drawbacks, like side-effects [20], habituation, dependence [21, 22], and impaired cognitive performance [23]. A non-pharmacological alternative aimed at enhancing overall sleep quality is cognitive behavioral therapy for insomnia [24]. However, this treatment is costly, time-consuming, and not widely accessible, nor does it directly address the lack of SWS. Therefore, there is an urgent need for novel, accessible interventions that specifically target SWS disturbances.

A recent and promising approach towards sleep enhancement involves non-invasive delivery of acoustic stimuli that are aligned with slow oscillations (SOs; ∼0.5-1.5 Hz), the hallmark of SWS [25,26]. Previous studies have demonstrated that such procedures can enhance SOs in the seconds immediately following acoustic stimulation, or during rhythmic stimulation at a frequency in the slow oscillation range (typically around 1 Hz) [25-33]. However, when using rhythmic stimuli, as most studies have done, the effects of stimulation appear to be self-limiting, fading out with each subsequent stimulus [34]. Possibly, the non-phase targeted stimuli in a rhythmic sequence interfere with the intrinsic oscillatory brain dynamics. In line with this notion, untargeted acoustic stimuli were shown to disrupting traveling slow waves [35].

An advanced and flexible method in the area of neuroelectrically-guided stimulation is modelling-based closed-loop neurostimulation (M-CLNS). M-CLNS models and predicts raw neurophysiological signals in real-time, allowing high stimulus targeting performance across a wide range of brain oscillations and oscillatory phases, including SOs [36]. M-CLNS has been used to precisely phase-lock single acoustic stimuli to the onset of slow oscillations, with particularly strong effects in enhancing SO activity both in the short and longer term, suggesting overall sleep deepening [37-39].

Until now, sleep enhancement using acoustic stimulation has predominantly been researched in laboratory settings using high-end EEG equipment. Such sleep laboratory settings are costly, staff-intensive and generally have limited capacity, restricting the number of participants that can be studied simultaneously. To further advance M-CLNS and potential clinical applications, there is a need for user-friendly EEG wearable devices that could accommodate larger numbers of subjects, over extended periods, both in clinical and home environments.

In this study, we assess whether M-CLNS with acoustic stimulation of SOs can be implemented using a self-applicable EEG headband. To ensure adequate signal quality and real-time access to raw data, we use the research-grade ZMax device (Hypnodyne Corp., Sofia, Bulgaria) [40, 41], which has been validated against PSG for signal quality [42]. The ZMax wearable records EEG signals from forehead derivations.

This electrode configuration offers several advantages for potential future consumer-or clinical-grade devices. First, bare-skin electrodes with smooth surfaces are generally more comfortable than ‘dry’ electrodes used in the hair, which often feature brush-like designs that press into the skin. Second, the forehead’s visibility allows users to position the EEG sensors correctly. Third, the forehead is a good location for detecting sleep slow oscillations, which are most prominently recorded in frontal regions. These factors support the feasibility of translating this approach to consumer or medical applications. Thus, this study represents an important first step toward the potential miniaturization of M-CLNS-based sleep enhancement using an EEG wearable device.

Specifically, this study will record the sleep EEG from the EEG headband and perform real-time modelling on this signal to predict oscillatory dynamics and target soft sound stimuli to the start of SO negative deflections (relative to derivations with mastoid or earlobe references). Healthy individuals will sleep while simultaneously wearing the EEG headband and gold-standard PSG equipment during a single night. The main aims of the study are to assess stimulus targeting performance over the headband and whether the acoustic stimulation will deepen sleep, as was observed in our previous polysomnography experiments. We analyze both short-term, stimulus-locked responses and more enduring effects across stimulated sleep and hypothesize that M-CLNS over the EEG headband will enhance SOs and delta activity over both time spans. The effects of stimulation on SWS dynamics are assessed using gold-standard PSG equipment, which provides the most reliable signals, while at the same time we assess the extent to which effects of stimulation can be observed in the signals of the EEG wearable. These validations of the wearable EEG relative to the gold-standard PSG could be particularly important in future contexts where the EEG wearable may be used as a standalone device.

## 2. Methods

### 2.1. Participants

Twenty-six healthy participants (14 males; mean ± SD age: 21.23 ± 2.76) participated in this study. None of the participants reported diagnosed sleep, neurological or psychiatric disorders or the use of prescription drugs known to affect sleep. Participants were asked to abstain from alcohol consumption and recreational drug use for 24 hours, and to avoid caffeine consumption for at least 7 hours preceding the experiment. Moreover, participants were asked to go to bed no later than 12:00 PM the night prior to the experiment, and to awaken no later than 08:00 AM the morning prior to the experiment. All participants provided written informed consent. This study was approved by the Ethics Committee at the University of Amsterdam.

### 2.2. EEG wearable data acquisition

EEG wearable data was acquired using the ZMax headband, positioned just above the eyebrows on the flattest available surface of the forehead. The ZMax headband was selected for this study due to it being one of the few EEG commercially available EEG wearables with real-time access to raw data [43]. Left- and right-frontal EEG signals, approximating F7-Fpz and F8-Fpz, were captured using Ag/AgCl semi-wet hydrogel electrodes.

In addition to EEG, 3-axis acceleration and photoplethysmography data was captured through sensors in the headband (this data was not used in this experiment). Data from all sensors was transmitted to a wireless receiver dongle, connected to a USB-hub located less than 1 meter from the participant’s pillow. The signals were sampled at 256 Hz with a bandwidth of 0.1-128 Hz and acquired through a Transmission Control Protocol socket. Data recording and analysis were conducted on a dedicated computer using a custom version of EventIDE (Okazolab Ltd, Delft, The Netherlands, v.21.11.2019), a visual programming software for designing and executing experiments using real-time and low latency signal processing.

It is important to note that the signals from the forehead derivation, in which a frontolateral electrode is referenced to a frontocentral one, are inverted in polarity compared to EEG derivations referenced to the earlobe or mastoid. Therefore, the findings should be interpreted as the results of targeting the SO negative deflection, where most often in CLNS research the SO positive deflection is targeted for stimulation [44]. The same is of course true for the concomitant PSG derivation.

### 2.3. PSG data acquisition

PSG data, consisting of 64-channel EEG, vertical- and horizontal-electrooculography, and chin electromyography were recorded using a Refa8 72-channel DC amplifier (TMSi, Oldenzaal, The Netherlands). EEG was captured using sized WaveGuard caps (ANT, Enschede, The Netherlands) with separate sintered AgCl reference electrodes placed on the earlobes (A1, A2). Signals were sampled at 512 Hz and recorded on a dedicated computer using a customized version of TMSi Polybench (TMS international, The Netherlands, v.1.34.1.4456). EEG impedances were verified to be below 10 kΩ at the onset of recording.

The data acquired from both the EEG wearable and PSG were time-synchronized using commonly recorded markers, created in EventIDE and sent to the PSG amplifier for simultaneous recording in the PSG data. Specific markers were recorded at the start and end of the sleep recording, as well as at the start and end of each experimental block and at the onset of each stimulus presentation.

### 2.4. M-CLNS algorithm

The M-CLNS algorithm, developed at our lab in collaboration with Okazolab Ltd (Delft, The Netherlands), was employed to continuously perform non-linear sine fitting on the most recent 1.6 seconds (calculus windows) of the EEG wearable raw F7-Fpz signal. Stimulus release was targeted at the 180° phase (i.e. halfway the negative deflection) within the forecasted signal in the SO frequency range of 0.5 – 1.5 Hz. Acoustic stimuli were released when the targeted phase was predicted to occur 24 – 34 ms in the future (search window), based on the best-fitting estimated model [36].

### 2.5. Stimulation protocol

Acoustic stimuli were presented using Creative GigaWorks T20 Series II stereo speakers and consisted of faded-in 100 ms pink noise pulses, calibrated to be 42.5dB(A) using a Rion NA-27 sound level meter. The calibration point, at approximately 50 cm distance from the midpoint between the left and right speaker (located ∼56 cm apart), was chosen to approximate participant head location during sleep.

PSG and EEG wearable signals were monitored in real-time by the experimenters. The M-CLNS SO detection algorithm was manually enabled at SWS onset. During the ensuing three-hour interval, acoustic stimuli were presented phase-locked to SOs, while stimulus presentation was manually paused whenever light sleep, REM sleep, movement or arousals were observed. Stimulus presentation occurred in 30-second blocks of either acoustic stimulation (STIM) or muted stimulation (SHAM). Stimulation blocks were alternated randomly and were interspaced with 30-second no stimulation blocks. The onset of presented stimuli was recorded in EventIDE and sent to the PSG system for simultaneous recording in Polybench.

### 2.6. Procedure

Participants slept a single night at the Amsterdam Sleep and Memory Lab. They arrived at 09:00 PM and were asked to perform a quick hearing test to confirm the acoustic stimulation’s sound level was within their hearing range. Participants went to bed between 11:00 and 11:30 PM and were given the opportunity to sleep a minimum of 8 hours, while simultaneously wearing both the EEG wearable and PSG equipment. No information was provided on the acoustic stimulation that was to occur during sleep.

### 2.7. Data analysis

Offline data was pre-processed using EEGlab 14.1.2b [45] in MATLAB R2016a. The PSG EEG data was down-sampled to 256 Hz, 0.1 -49 Hz band-pass filtered, and re-referenced to the average of A1 and A2. The same band-pass filter was applied to the EEG wearable data.

The PSG F7-Fpz signal was extracted for comparison to the concomitant EEG wearable derivation, to assess consistency and possible differences between the two signals. Sleep recording quality was evaluated through visual inspection of the signals and spectrograms (0.5 -48 Hz), which were created using the Chronux 2.11 Spectral Analysis toolbox [46] in MATLAB. Signal stretches containing artifacts were removed from both PSG and EEG wearable data before further analyses were performed.

Stimulus presentation blocks from F7-Fpz for both EEG wearable and PSG data were collapsed into STIM and SHAM data sets for each participant. These data sets were then exported as EDF files using EEGlab. The EDF files were further processed and analyzed in Python (3.11.9) using the Pandas (2.2.3), NumPy (2.1.3), MNE (1.9.0), SciPy (1.15.2), PyCircStat (0.0.2), Astropy (7.0.1), Statsmodels (0.14.4), and Matplotlib (3.10.0) libraries [47-55].

First, stimulus presentation accuracy for the STIM and SHAM conditions were calculated. To this purpose the data sets, containing stimulus onset markers, were band-pass filtered (0.5-1.5 Hz FIR; MNE), and the analytic signal was obtained via Hilbert transform (SciPy). The instantaneous phase was derived at each stimulus marker and a correction was applied by subtracting π/2 to account for the phase shift. Subsequently, the phase offset from the targeted phase was calculated in degrees for each corrected instantaneous phase. Circular statistics (mean, median, and standard deviation; Astropy and PyCircStat) were computed across STIM and SHAM for both EEG wearable F7-Fpz and PSG F7-Fpz to assess accuracy (mean/median phase offsets) and precision (phase offset standard deviations) of stimulus presentations [36].

To assess stimulus-locked effects, event-related potentials (ERP; MNE) were analyzed, again for EEG wearable F7-Fpz and PSG F7-Fpz. All STIM and SHAM data sets were band-pass filtered (0.5-1.5 Hz FIR; MNE), and epochs were extracted from -1 to 2 seconds around the stimulus markers. A baseline correction was applied using the pre-stimulus interval (−1 to 0 seconds). Grand average ERPs across all participants were then calculated for both the STIM and SHAM conditions. Paired samples t-tests (SciPy) were performed for each timepoint after stimulus onset, comparing STIM and SHAM. To account for multiple comparisons, a false discovery rate correction (Benjamini-Hochberg method; from the Python library Statsmodels) was applied, with a significance threshold of .05. Results were presented as smoothened bar plots to indicate significant differences between grand average ERPs.

Overall effects of M-CLNS with acoustic stimulation were assessed through power spectral density (PSD), analyzed for each participant and signal (PSG F7-Fpz and EEG wearable F7-Fpz) for both STIM and SHAM condition concatenated blocks. PSD was estimated using Welch’s method (SciPy), with power calculated in 0.25 Hz bins across the frequency range of 0.25-35 Hz. These bins were chosen to provide a fine-grained resolution of power, allowing for the detection of subtle differences in oscillatory activity. Normality of the data was assessed through Shapiro-Wilk tests and visual inspection of Q-Q plots. Participants whose data exhibited extreme outlier values as identified in Q-Q plots were excluded from further analysis for both PSG and EEG wearable. To compare power between the STIM and SHAM conditions, we conducted paired samples t-tests (SciPy) when the data met normality assumptions, and Wilcoxon signed-rank tests (SciPy) when the data did not meet normality assumptions, for each frequency bin across all participants. To control for multiple comparisons, we applied a false discovery rate correction (Benjamini-Hochberg; Statsmodels) with a significance threshold of .05. Additionally, we calculated the percentage difference in power between STIM and SHAM for each frequency bin to further illustrate power changes between conditions.

## 3. Results

### 3.1. EEG wearable signal quality

We evaluated and compared the quality of the EEG wearable F7-Fpz to PSG F7-Fpz signals, to assess the reliability of the EEG wearable. Out of the 26 EEG wearable recordings, nine showed no sign of data quality issues throughout the entire night. Another nine recordings showed incidental reduction of data quality, which only occurred after the stimulation interval (29 periods in total; M = 3.22, SD = 2.64). Six recordings exhibited incidental reductions in data quality both during (12 periods in total, M = 2, SD = 1.55) and after (15 periods in total, M = 2.5, SD = 1.52) the stimulation interval. Two recordings showed reduced data quality for more than 98% of the recording. In contrast, no data quality issues were observed in any of the 26 simultaneous PSG signals.

For participants with data quality issues during the stimulation interval, three were excluded from further analyses due to noisy stretches that covered the majority of the interval. Additionally, both participants with data issues for more than 98% of the recording were also excluded. Furthermore, five participants were excluded due to not completing the sleep interval (1), not showing signs of SWS (1), showing signs of arousal immediately following stimulus presentation (2), and computer issues (1). As a result, data from the remaining 16 participants were used for further analysis.

Visual inspection of EEG wearable and PSG signals demonstrated a high level of concordance between the two signals during SWS. However, the EEG wearable consistently exhibited lower overall amplitudes compared to the PSG F7-Fpz signals. It should furthermore be noted that, when comparing the EEG wearable and PSG F7-Fpz signals to frontal PSG signals using A1 and A2 as reference electrodes, the EEG wearable and PSG F7-Fpz signals are inverted in polarity.

Another difference between the EEG wearable and the concomitant PSG signals, is that the EEG wearable displayed a higher density of eye movement artifacts. This is likely due to the closer proximity of the EEG wearable sensors to the eyes. Furthermore, visual inspection of the EEG wearable signals revealed a greater presence of beta (13-30 Hz) frequencies and higher, which may be attributed to muscle activity (15 Hz and above) [56] that is recorded due to the closer proximity of the EEG wearable sensors to the frontal belly of the occipitofrontalis muscle and other facial musculature [57]. Finally, sweat artifacts were more prominent in the EEG wearable signals compared to the PSG signals, which may be explained by the differences in sensor types (patches with solid gel contacts on the EEG wearable versus gelled cups with a sintered electrode on the PSG cap) [58].

### 3.2. M-CLNS adequately targeted SO phase

SO targeting over the headband was shown to have occurred adequately based on stimulus onset markers recorded in the gold standard PSG derivation (F7-Fpz) adjacent to the EEG wearable derivation. Analyses of the EEG wearable F7-Fpz signals demonstrated similar targeting accuracy and precision.

A total of 2665 and 2558 stimuli were presented during the STIM and SHAM blocks, respectively. On average, 6.07 and 5.99 stimuli were presented per STIM and SHAM block. The phase offsets for the STIM condition were -25.47° ± 64.41° (mean ± SD) and -28.68° ± 59.75°, as recorded with PSG and EEG wearable, respectively (see figure 1). Similarly, the phase offsets for the SHAM condition were -30.09° ± 67.02° and -31.94° ± 62.78° for PSG and EEG wearable (see figure 1).

**Figure 1.**
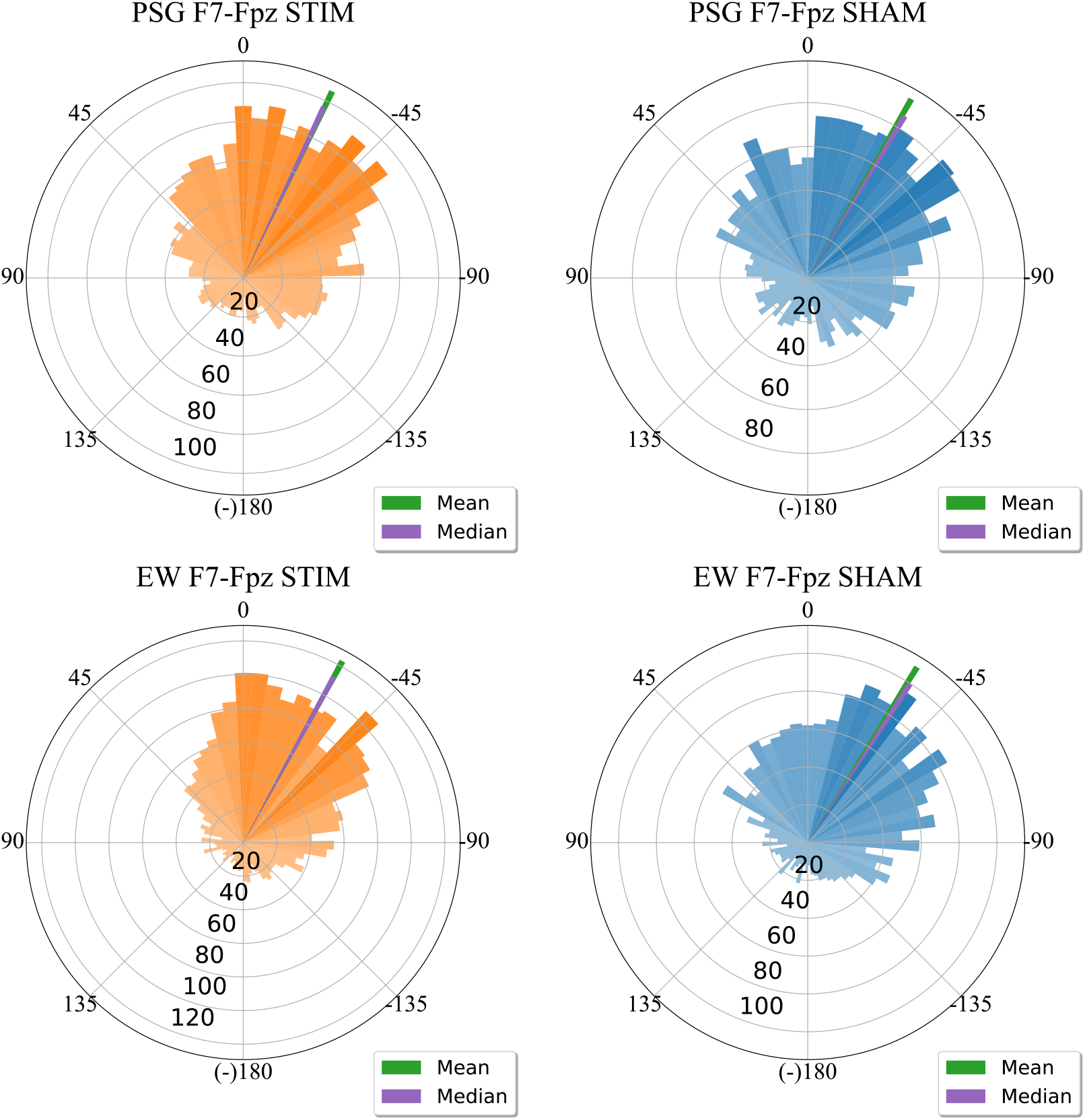
Phase targeting performance for slow oscillations recorded from a forehead derivation using an EEG wearable (EW). Circular histograms display the distribution of binned stimulus marker offsets (the difference between stimulus and target phase in degrees) for STIM (orange) and SHAM (blue) conditions, measured in both PSG F7-Fpz and EEG wearable F7-Fpz. The mean and median stimulus marker offsets are indicated by the green and purple lines, respectively.

These findings demonstrate that M-CLNS can target SO phase with accuracy and precision in forehead EEG wearable signals and that phase targeting accuracy can be assessed reliably from the EEG wearable signals.

### 3.3. M-CLNS with acoustic stimulation enhances slow oscillations in the short-term

Grand average ERPs reveal clear enhancement of a SO-like dynamic in the STIM compared to the SHAM condition, in both the PSG F7-Fpz and EEG wearable F7-Fpz signals. In both signals statistical comparison reveals three significant (p < .05) time ranges, reflecting a sequence of more pronounced slow positive and negative deflections in STIM compared to SHAM (figure 2). On average, these significant time ranges overlapped by 70.43% ± 26.54% between the grand average ERP findings for EEG wearable and PSG (table 1).

**Table 1.** Significant time ranges (p < .05; in ms) where M-CLNS with acoustic stimulation enhanced SO activity for both PSG F7-Fpz and EEG wearable F7-Fpz signals, along with the percentage overlap of these time ranges between the two signals.

| PSG F7-Fpz |  | EEG wearable F7-Fpz |  | EEG wearable-PSG overlap percentage (%) |
| --- | --- | --- | --- | --- |
| Start Time (ms) | End Time (ms) | Start Time (ms) | End Time (ms) |  |
| 351 | 648 | 113 | 723 | 48.69 |
| 801 | 1168 | 855 | 1117 | 100 |
| 1418 | 1746 | 1398 | 1922 | 62.60 |

**Figure 2.**
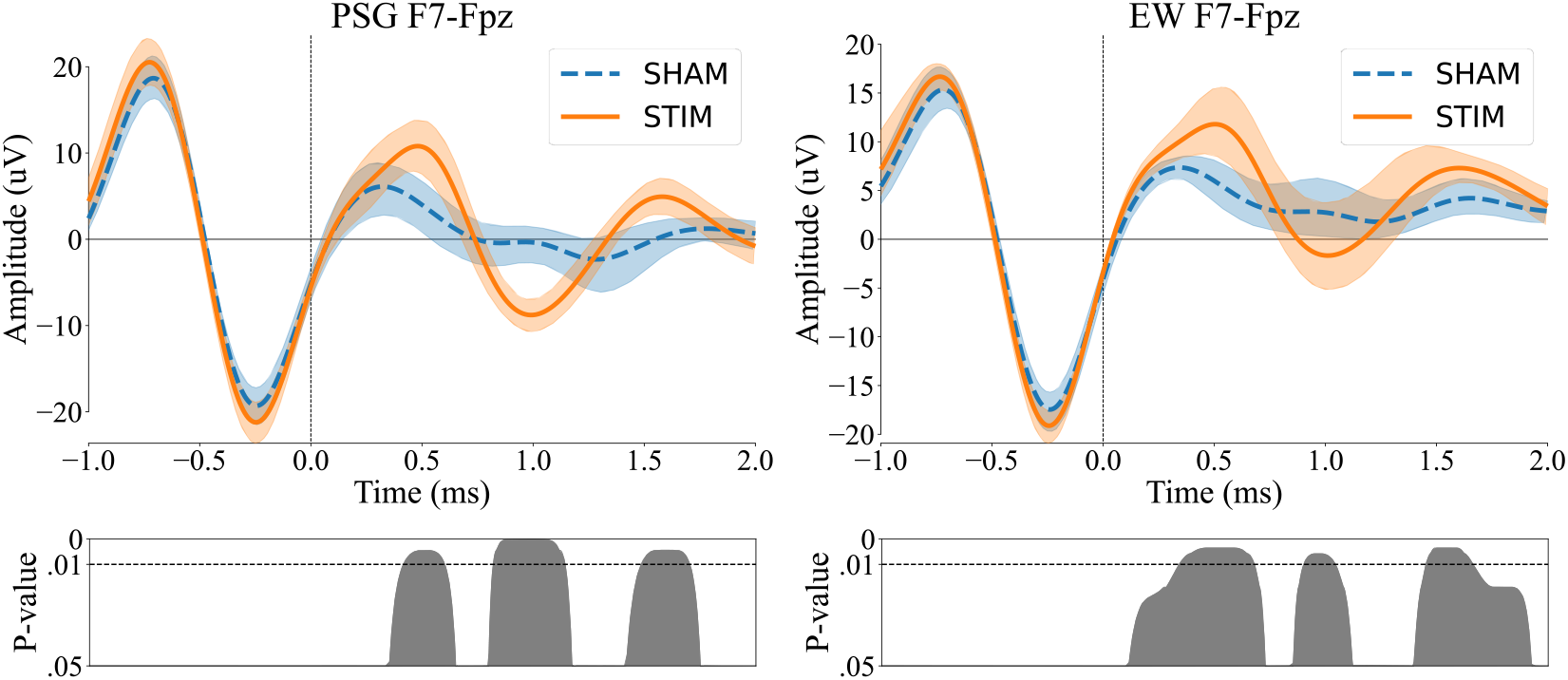
Stimulus-locked effects of M-CLNS with acoustic stimulation targeting SOs in EEG wearable (EW) signals and simultaneous PSG. Grand average ERP plots are presented comparing STIM and SHAM conditions for both PSG F7-Fpz and EEG wearable F7-Fpz signals, with corresponding 95% confidence intervals. Significant differences between STIM and SHAM, based on FDR-corrected paired samples t-tests, are indicated in accompanying plots, in which he significance threshold of p < .01 is marked with a dotted line.

These results highlight a reliable and substantial immediate enhancement of SO-like deflection in both signals. This suggest that M-CLNS with acoustic stimulation effectively modulates SO activity, with the observed effects being both strong and consistent across PSG and EEG wearable signals.

### 3.4. M-CLNS with acoustic stimulation leads to overall sleep deepening

To assess overall effects of M-CLNS with acoustic stimulation on the sleep EEG’s frequency content, we compared power spectral density between STIM and SHAM in 0.25 Hz bins across the 0.25-35 Hz frequency range. PSD data from two participants were excluded from further analysis due to exhibiting extreme outlier values as identified in Q-Q plots, resulting in a final sample size of N=14. The comparisons revealed significant power increases for STIM versus SHAM from 0.75 to 2 Hz, at 2.75 Hz, and from 3.25 to 4 Hz in the PSG signals. EEG wearable signals showed similar power increases from 0.75 to 5 Hz (table 2; figure 3).

**Table 2.** Means, standard deviations, paired samples t-test statistics, and percentage difference comparing frequency bin Welch spectral power between STIM and SHAM conditions for both PSG F7-Fpz and EEG wearable F7-Fpz signals. Only frequency bins showing statistically significant differences are included in the table.

| Hz | PSG F7-Fpz |  |  |  |  |  |  | EEG wearable F7-Fpz |  |  |  |  |  |  |
| --- | --- | --- | --- | --- | --- | --- | --- | --- | --- | --- | --- | --- | --- | --- |
|  | STIM |  | SHAM |  | Diff |  |  | STIM |  | SHAM |  | Diff |  |  |
|  | Mean | SD | Mean | SD | (%) | *t*(13)/*W* | *p* | Mean | SD | Mean | SD | (%) | *t*(13)/*W* | *p* |
| 0.75 | 196.25 | 91.26 | 166.76 | 74.61 | 17.68 | 4.25 | .021 | 129.71 | 49.64 | 108.37 | 49.82 | 19.69 | 5.59 | .006 |
| 1.00 | 215.27 | 105.17 | 170.85 | 81.21 | 26.00 | 5.10 | .021 | 110.85 | 32.38 | 88.87 | 27.22 | 24.73 | 6.55 | .003 |
| 1.25 | 172.10 | 91.83 | 137.04 | 70.69 | 25.58 | 4.52 | .021 | 83.45 | 24.90 | 68.84 | 20.02 | 21.22 | 1.00 | .007 |
| 1.50 | 110.92 | 57.48 | 88.24 | 44.29 | 25.70 | 4.50 | .021 | 55.42 | 18.28 | 45.47 | 14.43 | 21.87 | 5.07 | .007 |
| 1.75 | 68.18 | 33.91 | 53.65 | 25.66 | 27.08 | 4.86 | .021 | 35.40 | 12.81 | 29.08 | 10.72 | 21.75 | 4.35 | .012 |
| 2.00 | 42.20 | 21.31 | 34.12 | 16.71 | 23.69 | 4.17 | .021 | 22.49 | 8.81 | 19.00 | 8.29 | 18.36 | 3.81 | .025 |
| 2.25 |  |  |  |  |  |  | ns | 14.99 | 6.24 | 12.97 | 6.08 | 15.58 | 3.53 | .032 |
| 2.50 |  |  |  |  |  |  | ns | 10.52 | 4.64 | 9.20 | 4.15 | 14.34 | 4.41 | .012 |
| 2.75 | 13.56 | 6.89 | 11.47 | 5.23 | 18.24 | 5.00 | .021 | 7.65 | 3.32 | 6.64 | 2.80 | 15.33 | 4.81 | .007 |
| 3.00 |  |  |  |  |  |  | ns | 5.84 | 2.27 | 5.06 | 2.04 | 15.48 | 4.30 | .012 |
| 3.25 | 8.14 | 3.64 | 7.12 | 3.10 | 14.29 | 3.66 | .040 | 4.67 | 1.63 | 4.11 | 1.59 | 13.64 | 3.96 | .021 |
| 3.50 | 6.62 | 2.87 | 5.91 | 2.51 | 11.91 | 4.15 | .021 | 3.86 | 1.31 | 3.41 | 1.26 | 13.05 | 5.35 | .006 |
| 3.75 | 5.67 | 2.56 | 4.97 | 2.09 | 14.00 | 3.62 | .040 | 3.30 | 1.12 | 2.89 | 1.05 | 14.40 | 4.91 | .007 |
| 4.00 | 5.02 | 2.29 | 4.32 | 1.88 | 16.16 | 3.68 | .040 | 2.85 | 0.97 | 2.48 | 0.94 | 14.92 | 3.66 | .029 |
| 4.25 |  |  |  |  |  |  | ns | 2.40 | 0.84 | 2.12 | 0.80 | 13.51 | 3.55 | .032 |
| 4.50 |  |  |  |  |  |  | ns | 1.99 | 0.69 | 1.79 | 0.62 | 11.33 | 3.68 | .029 |
| 4.75 |  |  |  |  |  |  | ns | 1.71 | 0.59 | 1.55 | 0.51 | 10.42 | 3.49 | .033 |
| 5.00 |  |  |  |  |  |  | ns | 1.52 | 0.52 | 1.37 | 0.45 | 10.61 | 3.46 | .033 |

PSG = polysomnography. ns = non-significant. Diff = percentage difference between STIM and SHAM. Positive difference scores indicate higher spectral power in STIM compared to SHAM.

**Figure 3.**
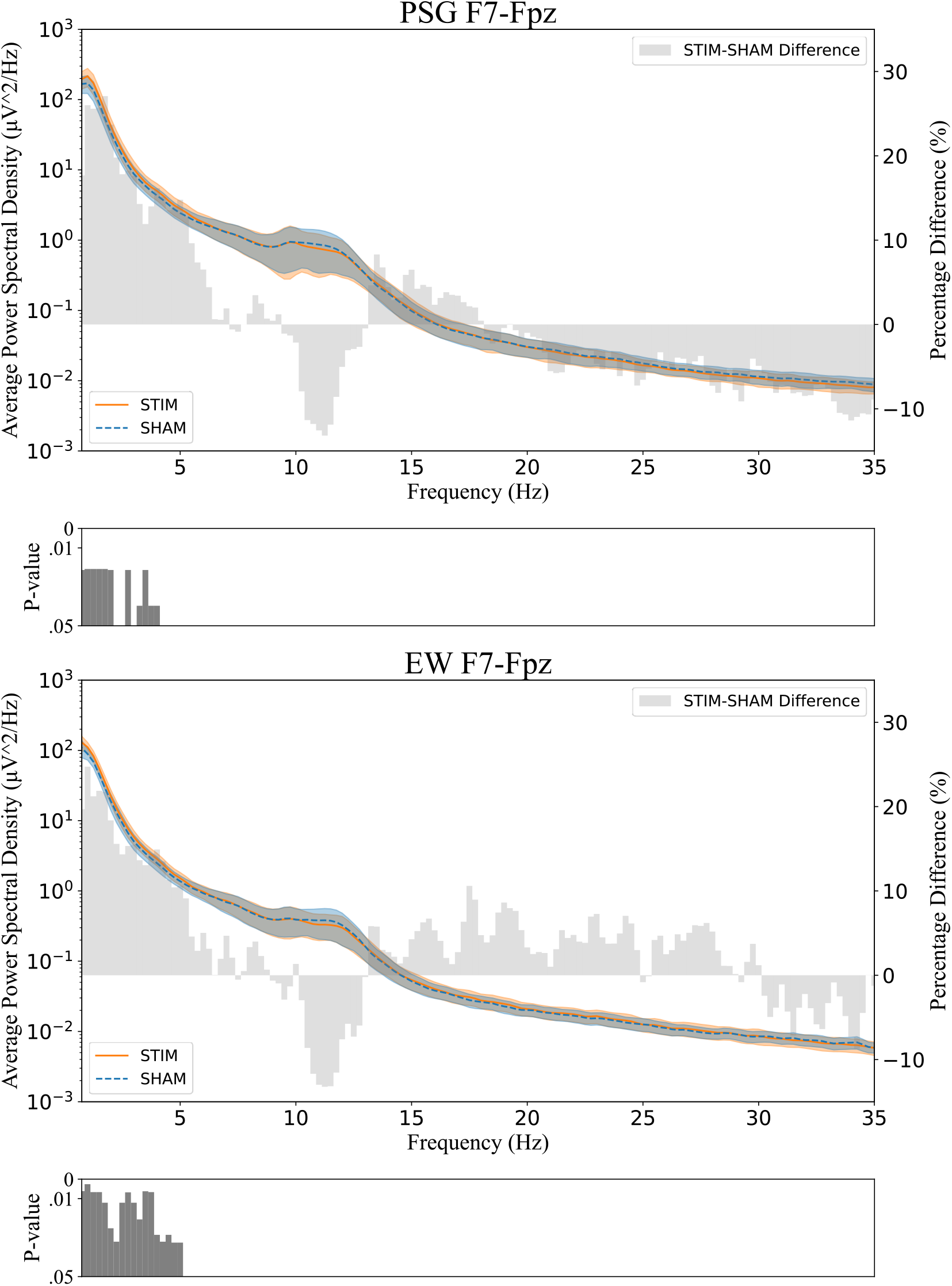
Power spectral density (PSD). PSD plots comparing STIM and SHAM conditions in 0.25 Hz bins across the 0.25-35 Hz frequency range for both PSG F7-Fpz and EEG wearable (EW) F7-Fpz signals. The plots include 95% confidence intervals and percentage differences in power between STIM and SHAM. Significant differences, based on FDR-corrected paired samples t-tests, are indicated in accompanying bar plots.

The average increase in power, based on the percentage difference between STIM and SHAM across significantly increased frequency bins in the delta range (0.75-4 Hz) was 20.03% ± 4.66% for PSG and 17.45% ± 3.74% for EEG wearable signals. Considering only the SO range (0.75-1.5 Hz), the average bin power increase across significantly increased frequency bins was 23.74% ± 4.04% and 21.88% ± 2.11% for PSG and EEG wearable data, respectively (table 2).

While no other statistically significant differences were found between the STIM and SHAM conditions in either device, two other observations merit attention. First, a non-significant, but notable decrease in power was observed in the low sigma range, from approximately 9 to 13 Hz, in both the PSG and EEG wearable signals (figure 3). This could reflect an effect on slow sleep spindles, which occur at this frequency. Secondly, the PSG show a trend for decreased power with stimulation in the beta range (13-30 Hz), while the EEG wearable shows a tendency for increased values. The latter observation might reflect increased muscle activity during stimulation, picked up by EEG wearable sensors due to their close proximity to facial musculature.

These findings show that M-CLNS using acoustic stimulation can enhance SO and delta activity across sleep. This power enhancement can be reliably picked up in the EEG wearable signals. The non-significant reduction in power within the slow spindles range might reflect a mild suppression of slow spindles. Finally, caution should be used in interpreting EEG wearable signals in the beta range, given possible contamination with muscle activity or other artifacts in this range.

## 4. Discussion

In this study we evaluated whether sleep could be deepened using M-CLNS on an EEG wearable with forehead electrode derivations. We found that single non-arousing sound pulses could be precisely phase-targeted to slow oscillations (SOs) based on the wearable’s signal and that stimulation evoked enhanced slow oscillation-like responses in stimulus-locked analyses. Moreover, stimulation led to overall sleep deepening, reflected in enhanced SO and delta power across stimulated sleep.

We, furthermore, demonstrated that the EEG wearable signal accurately reflects the effects of acoustic stimulation in the form of enhanced power in the SO and delta range. This despite differences in the EEG wearable signal relative to PSG, such as reduced amplitudes, more prevalent artifacts (due to muscle activity, eye movements, and sweat) and incidental signal loss. The short-term, stimulus-locked effects of stimulation could also be clearly observed in the wearable data. In fact, both the short and long-term effects were similar between gold-standard PSG F7-Fpz and EEG wearable F7-Fpz signals. Of note, we observed a non-significant but notable power reduction in the low sigma range, in which slow spindles occur.

The above findings show that, in principle, sleep deepening with M-CLNS and a read-out of the method’s effectiveness can be achieved using an EEG wearable with forehead derivations. Besides advancing the application of M-CLNS towards wearable devices, the findings corroborate and extend our previous results, which demonstrated overall sleep deepening using M-CLNS on PSG recordings during a nap [38]. The current study shows that overall sleep deepening also holds when SO stimulation is extended across an entire night of sleep. In line with this previous nap study, a similar, non-significant but notable, reduction of sigma power was observed [38]. These trends might be related to the inverse relation between NREM sleep depth and spindle genesis and are thus in line with an overall sleep deepening through M-CLNS [59].

Sleep enhancement using EEG wearables has also been attempted using other EEG wearables, using more classical frontal to mastoid derivations, and different stimulation methods. Many of these studies have used rhythmic stimulation, over the SmartSleep Deep Sleep Headband (Philips Healthcare, the Netherlands) [60-63], or stimulation trains with the DREEM headband (Rythm SAS, Paris, France) [64-65]. These studies assessed effects during rhythmic stimulation [60-63] or short-term effects in a few seconds after onset of individual stimuli [64-65]. Overall, these studies find the well-known immediate effects of enhanced SO-like dynamics in ERP analyses together with increases of stimulus-locked delta power [64-65], or increases of delta power as long as rhythmic stimulation is ongoing [60-63]. In contrast to the current study, these studies have not demonstrated enduring effects of single SO phase-targeted stimuli across SWS, suggesting only an enhancement of SOs and delta power in a few seconds after stimulus onset, or during rhythmic stimulus trains. As found with similar protocols run over gold standard equipment, these effects of “driving stimulation” appear to be self-limiting [34].

Studies using a protocol with single SO phase targeted stimuli, implemented on versions of the Mobile Health Systems Lab Sleep Band (MHSL-SB; ETH Zurich, Zurich, Switzerland) [66] have demonstrated increases of low slow-wave activity (0.75-1.25 Hz) when comparing between stimulation and sham nights, both in healthy adults [67] and recently in a proof-of-concept study involving Parkinson’s patients [68]. A study involving healthy young adults found a broader range of increases of delta-theta and sigma power when comparing stimulation to sham nights [69].

Interestingly, SO stimulation may also be feasible using in-ear EEG devices [70]. Although the correlation between in-ear (or across-ear) EEG signals and frontal scalp signals is significantly lower than that of a forehead derivation, the potential comfort offered by in-ear EEG devices specifically designed for sleep makes this form factor worth exploring.

A few points regarding the use of sleep enhancing algorithms over EEG wearables merit consideration. First, while in this study delta power enhancements could be reliably measured over the wearable, analysis of phenomena in the beta range should be undertaken with caution. Increased power in this range compared to a PSG derivation adjacent to the wearable suggests the beta range might be polluted with possible muscle activity.

More in general it is worth noting that, for EEG self-applicable devices, signal quality and consistency across the night, will vary with the type of electrodes, the derivation used for recording, the form factor, and with other aspects of the wearable design. A device like the MHSL-SB remains relatively close to gold-standard equipment, using pre-gelled silver/silver chloride (or even gold) electrodes, and recording from Fpz referenced to M2 with M1 as the ground electrode [66]. According to the developers, the signal is similar to that of a CE-certified PSG device, with only a small amplitude reduction [66]. In a device like ZMax, which records from a forehead derivation, and thus captures more local potential differences, the amplitude of the signal is considerably reduced compared to a frontal to mastoid derivation (by about 40%) [42]. In-ear EEG devices have an even lower amplitude profile and a lesser correlation with PSG frontal to mastoid derivations. Consumer-grade EEG wearables, which typically feature so-called ‘dry’ electrodes intended for long term use, will tend to have lower signal quality and signal consistency compared to research and medical grade devices, which feature single use or short-term use electrodes. At home sleep recording and stimulation studies with consumer grade devices may therefore be challenging, as exemplified by a study in which the DREEM device was used to stimulate SOs in patients with Alzheimer’s disease over a 14-day period [65]. In this study overall stimulation frequency was low, with 57% of the recorded nights not featuring any stimulation at all [65].

While EEG wearables present challenges, another issue concerns SO stimulation protocols. Most methods adopted thus far (signal amplitude-based, PLL procedures, phase vocoder) tend to function best on high amplitude SOs but struggle to target lower amplitude waves [71]. This presents a drawback in the context of populations with reduced SO dynamics, including the elderly and most disorders associated to sleep problems. It also limits applications on EEG wearables with, for instance, forehead or in-ear signal recording, which intrinsically feature reduced amplitudes with regards to standard PSG derivations. Here our M-CLNS approach offers an advantage, as the pertaining algorithm is not intrinsically dependent on, nor sensitive to, signal amplitude.

The results of this study should be interpreted within the context of a few limitations. First, the study consisted of healthy young individuals. While this population was purposely selected for this first transition to a wearable approach, the findings should not be generalized to older or diseased populations.

Additionally, the single night, within subject design of this experiment means that, in theory, sleep deepening in stimulation blocks could lead to some compensatory lightening of sleep in buffer and sham stimulated blocks. Hence, overall sleep deepening effects of M-CLNS stimulation needs to be confirmed in a mixed multi-night design. In study, though using a different acoustic stimulation algorithm, it is suggested that sleep deepening effects of SO phase targeted single stimuli can persist across multiple days of sleep stimulation [67].

Finally, it should be noted that the current findings were based on measurements in a laboratory setting. While this setting was intentionally chosen in this first feasibility study to also assess findings over gold-standard PSG, the application in the home-environment may come with additional challenges. Potential influencing factors include environmental noise, poor bedroom sleep hygiene or improper application of wearable devices, which could negatively affect sleep or signal quality. Future studies transitioning to the home environment should carefully consider such potential confounding factors.

The current M-CLNS sleep deepening effects may provide valuable insights for populations experiencing a decline in SWS. An example hereof is the globally increasing proportion of the ageing population, in which SWS typically decreases [72]. While there may be a growing need to address SWS deficits, especially given the ongoing health epidemic of sleep problems as declared by the WHO, current treatments generally do not specifically address this issue. M-CLNS may offer a novel approach that directly targets enhancing SWS.

Additionally, M-CLNS may offer a promising strategy for a wide range of clinical populations that suffer from a lack of SWS. These include patients with insomnia [11,12], sleep apnea [13] and parasomnias, as well as psychiatric conditions like post-traumatic stress disorder [73] and depression [15]. Somatic diseases like chronic fatigue syndrome and long-COVID may also be targeted, given their reductions in SWS [74]. Importantly, restoring SWS sleep may help to delay or even prevent neurodegenerative diseases, including Alzheimer’s [16] and Parkinson’s disease [17]. A wearable application may serve as a valuable tool in clinical inpatient settings, offering a non-pharmacological approach to restore SWS across various populations and treatment environments.

## 5. Conclusion

In conclusion, we have demonstrated that M-CLNS with acoustic stimulation can be utilized, using a forehead EEG wearable to effectively target SOs resulting in immediate, as well as enduring enhancement of slow oscillations. M-CLNS may offer a promising, portable and non-pharmacological tool to enhance SWS, with future potential for application in the home environment and clinical settings.

## Data availability statement

The data that support the findings of this study are available upon reasonable request from the authors.

## Author contributions

Conceptualization & Methodology: S L J and L M T; Data collection: S L J; Programming: S L J and A C V D H; Formal Analysis: S L J; Supervision: L M T; Writing: all authors contributed to manuscript writing and editing; All authors have read and agreed to the published version of the manuscript.

